# Gametocyte production and transmission fitness of African and Asian *Plasmodium falciparum* isolates with differential susceptibility to artemisinins

**DOI:** 10.1101/2024.12.23.630038

**Authors:** Nicholas I. Proellochs, Chiara Andolina, Jordache Ramjith, Rianne Stoter, Geert-Jan van Gemert, Wouter Graumans, Susana Campino, Leen N. Vanheer, Martin Okitwi, Patrick K. Tumwebaze, Melissa D. Conrad, Taane G. Clark, David A. Fidock, Didier Menard, Sachel Mok, Teun Bousema

## Abstract

The emergence of *Plasmodium falciparum* parasites partially resistant to artemisinins (ART-R) poses a significant threat to recent gains in malaria control. ART-R has been associated with PfKelch13 (K13) mutations, which differ in fitness costs. This study investigates the gametocyte production and transmission fitness of African and Asian *P. falciparum* isolates with different K13 genotypes across multiple mosquito species. We tested three ART-sensitive (ART-S) isolates (NF54, NF135, NF180) and three ART-R isolates (ARN1G, 3815, PAT-023) for sexual conversion and transmission to *Anopheles stephensi, An. gambiae* and *An. coluzzii*. ART-R levels were quantified *in vitro* using the Ring-stage Survival Assay (RSA), and the transmission-reducing effects of dihydroartemisinin (DHA) on mature gametocytes were assessed. Results showed that ART-S parasite lines consistently produced gametocytes and transmitted effectively in all three mosquito species. ART-R isolates showed variability: ARN1G maintained high transmission levels, whereas 3815 showed limited transmission potential despite higher sporozoite loads in *An. coluzzii*. The African ART-R isolate PAT-023 demonstrated low gametocyte commitment but was transmitted efficiently in both *An. gambiae* and *An. coluzzii*. DHA exposure reduced mosquito infectivity for all isolates, regardless of K13 genotype. These findings, based on a limited number of field isolates, suggest that ART-R parasites remain transmissible across different *Anopheles* species. However, ART-R does not appear to confer a direct transmission advantage. This study highlights the complexity of ART-R dynamics and underscores the need for further research to inform malaria control strategies in regions where ART-R parasites are circulating.

## Introduction

Despite global efforts to reduce malaria burden, progress has plateaued in the last decade and in some areas malaria is again increasing [1]. The recent emergence in Sub-Saharan Africa of *P. falciparum* parasites with partial resistance to artemisinins (ART-R) adds to the concerns about how sustainable malaria control may be. ART-R is characterised by prolonged parasite clearance times (half-life >5 hours) or persistence of parasitemia on day 3 following artemisinin monotherapy or artemisinin-based combination therapy (ACT) [2–6]. ART-R is predominantly associated with specific mutations in the essential PfKelch13 (K13) protein’s β-propeller domain [7–10]. These mutations reduce K13 protein levels, disrupting haemoglobin import into the parasite’s digestive vacuole, a critical step for ART activation [8, 11–13]. While there are validated K13 mutations that associate with ART-R, their impacts on resistance and parasite fitness are variable and highly dependent on the mutation and parasite background [13–16]; as such it is difficult to predict which mutations are of particular concern.

Clinically relevant mutations in K13 first were observed in isolates collected in 2002 in western Cambodia [17]. While partial resistance subsequently spread across the Greater Mekong Subregion [7, 18], the anticipated migration to the African continent has not occurred. It was hypothesised that lower drug pressures combined with high parasite diversity in many settings in Sub-Saharan Africa may have allowed wild-type parasites to outcompete less-fit K13 mutant parasites, making *de novo* emergence of ART-R less likely and limiting the spread of the K13 mutants upon introduction. The recent independent emergence of K13 mutations associated with ART-R including in East Africa (Uganda, Rwanda, and Tanzania) and the Horn of Africa (Eritrea, Ethiopia and Sudan) [19–23] raises questions about their fitness, resistance, and transmission potential. Transmission to mosquitoes depends on the formation of viable male and female gametocytes, with considerable variation in gametocyte production between parasite isolates [24, 25]. The transmission potential of parasites with K13 mutations is particularly relevant with changes in vector populations and variation in *P. falciparum* vector competence [26]. The recent invasion of the competent Asian vector *Anopheles stephensi* in urban African settings [27, 28] raises concerns about the spread of ART-R parasites of African and Asian origin in settings previously not endemic for malaria.

Understanding variations in vector competence and determining whether *K13* mutant parasites possess a transmission advantage, both in the presence and absence of artemisinin derivatives, are crucial for predicting the potential spread of ART-R on the African continent. In this study, we investigate the gametocyte production and transmission fitness of ART-S and ART-R field isolates. We compared ART-S isolates [29, 30] with ART-R isolates carrying two common Southeast Asian K13 mutations G449A and C580Y [18, 31, 32], as well as a recently isolated ART-R K13 independent parasite line from Uganda, where the local emergence of ART-R has been documented including through K13 mutations.

## Material and methods

### Parasite cultures

All parasite lines – including the ART-S NF54 (Africa), NF135 (Cambodia) and NF180 (Uganda), as well as the ART-R *P. falciparum* field isolates ARN1G (Thailand), 3815 (Cambodia), and PAT-023 (Uganda) – were cultured in an automated culture system using RPMI media supplemented with 10% human serum, with gametocyte cultures for mosquito feeding set up at a uniform 1% parasitemia and allowed to mature for 14 days prior to feeding [33]. Asexual parasite cultures were maintained for no more than 25 cycles in a synchronised state through magnet separation or sorbitol lysis. Gametocyte induction was performed using either minimal fatty acid media or 0.5% Albumax; gametocytes were allowed to mature in RPMI media supplemented with 10% human serum [25]. Parasite DNA was extracted with QIAGEN Blood DNA kit, sequenced on the Illumina Novaseq 6000 platform and analysed using the malaria profiler tool [34].

### Ring-stage Survival Assay (RSA)

The Ring-stage Survival Assay (RSA) was performed as previously published [16, 35] with minor modifications. To avoid the negative impact of sorbitol synchronisation on conversion rates within the same cycle [36, 37], we opted for a gentler approach using double magnet purification. Highly synchronised parasites were first passed through a magnetic column (MACS) to collect segmented schizonts. These schizonts were put back into culture for 3-4 hours to allow bursting and reinvasion of new red blood cells (RBCs). The culture was subsequently passed through the same magnetic column, isolating the flow-through, which contained only newly invaded ring-stage parasites. Parasites were split into 2 plates: one for the sexual conversion assay and one for the RSA. For the RSA, parasites were diluted to 1-2% parasitemia in 2% hematocrit. Each line was tested in 4 wells: two for DMSO controls and two for DHA. Washes were performed in separate tubes following DHA treatment before being added to a fresh plate. Survival rates were determined using both Giemsa smears and flow cytometry using MitoTracker Red as a live cell stain from the same well for all replicates.

### Sexual conversion assay

Sexual conversion rates were performed as previously published [25], with minor modifications. After double synchronisation, parasites were returned to culture for 24 hours prior to initiating the conversion assay. The assay was performed in a plate-based format with each parasite isolate split over 3 wells (one for each media type) at 1% parasitemia in 5% hematocrit. Conversion rates were calculated by Giemsa-stained smears [25].

### Mosquito infections

Laboratory colonies of *An. stephensi* (Nijmegen Sind-Kasur strain)[38], *An. coluzzii* (N’gousso strain) [39] and *An. gambiae* s.s (Kisumu strain) [40], were maintained under controlled conditions: 26°C, 70-80% humidity and a 12-hour reverse day/night cycle. Gametocyte cultures were checked for exflagellation to confirm maturation and gametocytaemia reached a minimum of 0.1% before feeding. Culture material was not further normalized for gametocyte density prior to feeding. Groups of 100 female *Anopheles stephensi*, *An. gambiae*, and *An. coluzzii* mosquitoes, aged 1–5 days, were blood-fed for 15 minutes using glass membrane mini-feeders (15 mm diameter, convex bottom) connected to a heated circulating water bath. Fully fed mosquitoes were selected, and maintained at 30°C with access to 5-10% glucose.

### Ookinetes count

Mosquito midguts were examined 18-24 hours post-infection to identify the presence of round forms, retort forms, and mature ookinetes. Five midguts from each infection group were dissected and incubated with a 1:50 diluted Anti-25KD-FITC conjugate in Evans blue solution. Midguts were gently disrupted using a pipette tip to release the blood meal. Following incubation in the dark for 30 minutes at room temperature, the solution was washed with 1.4 mL of phosphate-buffered saline (PBS), vortexed to dissolve the pellet, and centrifuged for 2 minutes at 10,000 rpm. After removal of the supernatant, the pellet was resuspended in 25 µL of PBS and 5 µL of the suspension was loaded into a Bürker-Turk counting chamber. Round forms, retort forms and mature ookinetes were counted using an incident light fluorescence microscope with a GFP filter at 400× magnification.

### Oocysts count

Seven days post-infection, 20 mosquito midguts per group were dissected, stained with 1% mercurochrome, and examined under an optical microscope at 100× magnification to detect and quantify oocysts.

### Sporozoites count

Mosquitoes used to examine sporozoite development received a second (uninfected) bloodmeal on day 4-6 post infection to synchronise oocyst development Fifteen days post-infection, mosquito salivary glands were dissected in PBS and transferred to oocyst lysis buffer (NaCl 0.1M: EDTA 25mM: TRIS-HCl 10mM). Following overnight incubation at 56°C with Proteinase K, DNA was extracted with the automated MagNA Pure LC instrument using the MagNA Pure LC DNA Isolation Kit – High performance. Sporozoite density was analysed using COX-I qPCR [41].

### Statistical analysis

Statistical analyses were conducted using R software (v 3.1.12) [42]. Mean counts of oocysts, sporozoites, and ookinetes (with 95% confidence intervals) were estimated using a mixed-effects negative binomial regression model, incorporating random intercepts for biological replicates and fixed effects for parasite line, mosquito species, and their interactions. Mean survival and commitment rates (with 95% confidence intervals) were calculated using a mixed-effects beta regression model. Transmission-reducing activity (TRA) and transmission-blocking activity (TBA)—reflecting reductions in oocyst density and the proportion of infected mosquitoes, respectively—were assessed for 700 nM and 7000 nM DHA using a Bayesian Poisson regression model.

## Results

### Parasite Backgrounds

We used well-characterised *P. falciparum* reference lines from Africa (NF54), Southeast Asia (NF135, Cambodia) [29], and a new isolate from East Africa (NF180, Uganda). These lines were complemented with previously published ART-R K13 mutant parasite lines from Thailand (ARN1G, K13-449A) [32], from Cambodia (3815, K13-580Y) [31, 35] and a recently collected isolate from Uganda with *in vitro* ART-R that is independent of K13 mutations (PAT-023). Genotyping revealed that some isolates harboured mutations associated with resistance to chloroquine and sulfadoxine-pyrimethamine (Table 1).

**Table 1.**
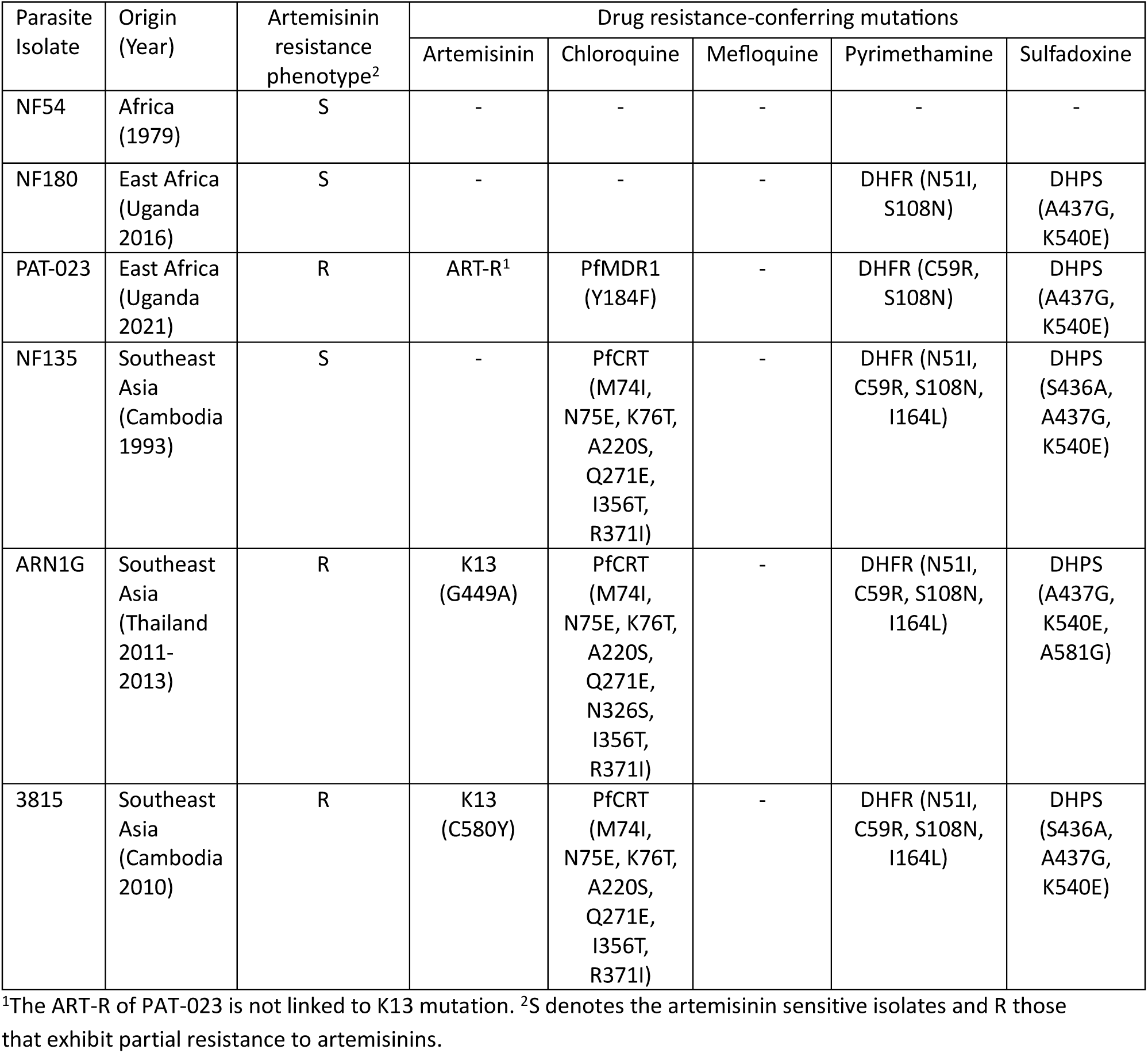
Parasite isolates.

### Artemisinin resistance

ART-R lines ARN1G and 3815 showed survival rates in the RSA that were higher than the ART-S NF54, as previously published ([16, 32, 35];Figure 1a, Sup Table 1). The ART-S NF135 line showed variable survival rates, which were lower than those observed for known ART-R lines but higher than those observed for ART-S NF54 and NF180 lines. The PAT-023 parasite line exhibited a survival rate of 9.1% (95% CI: 6.6% - 12.5%), higher than the ART-S lines.

**Figure 1.**
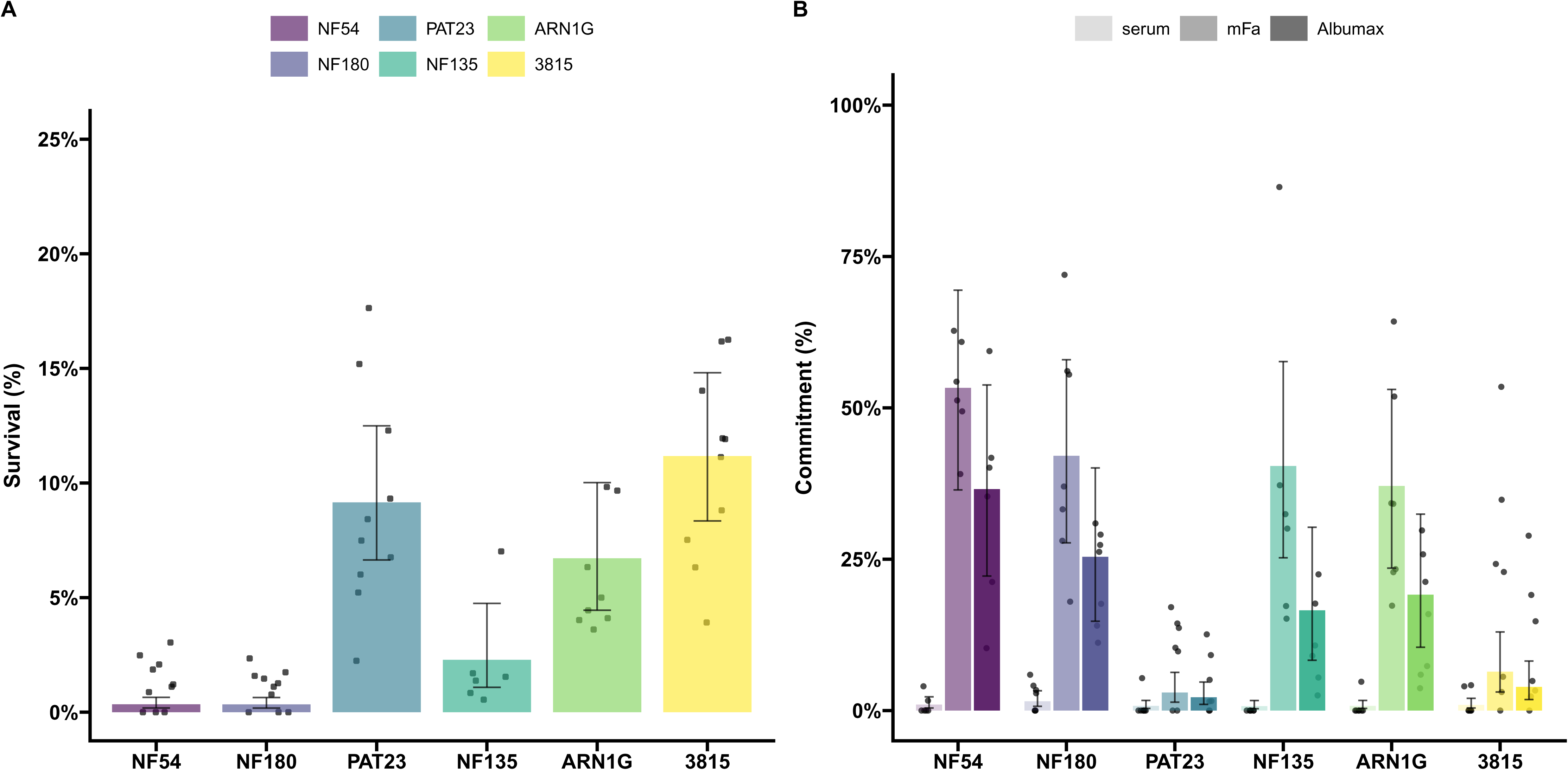
Parasite survival after exposure to dihydroartemisinin (DHA) and gametocyte conversion rates of *P. falciparum* isolates with different K13 genotypes and genetic backgrounds. **A)** Survival rate from the Ring-stage Survival Assay (RSA) of each parasite line following 6hr exposure to 700nM DHA compared to DMSO controls in duplicate wells (4-6 replicates were performed for each parasite line). The dots represent biological replicate survival rates at 72h post invasion from counts either by microscopy or flow cytometry and the error bars represent the 95% confidence intervals. **B)** Sexual conversion in a plate-based assay that quantifies conversion rates from a single asexual round using three different media types (shown different colour gradient of the bars). Conversion rates are calculated by dividing the final gametocytaemia by the starting parasitemia in the same well. Dots represent a single well from independent plates; error bars represent 95% confidence intervals.

### Sexual conversion

We assessed the ability of ART-R and ART-S lines to convert under two different conditions that mimic the natural sensing system [25, 43, 44]. All conversion assays were performed on a single asexual cycle. The highest rates of gametocyte conversion were typically observed using minimal fatty acid media with intermediate conversion rates for media containing Albumax only, and lowest conversion in non-inducing serum media (Figure 1b, Sup Table 2) [25]. The conversion rates observed for K13 mutants were comparable to those recorded for ART-S lines, except for the Cambodian line 3815, which exhibited highly variable conversion rates (Figure 1b). The Ugandan ART-R line PAT-023 exhibited minimal sexual conversion with little difference between media types (Figure 1b, Sup Table 2).

### Transmission to mosquitoes

Transmission was examined using three mosquito vector species. We first assessed whether fertilisation occurred in mosquitoes for each parasite isolate by quantifying parasite developmental stages in the mosquito midgut 20-24 hours post-infectious blood meal. We distinguished between mature and immature stages following Pfs25 antibody labelling. Immature stages included underdeveloped ookinetes and rounded forms classified as either zygotes or unfertilised females. The ART-S parasite isolates NF180 and NF135 showed no clear difference in the numbers of immature or mature ookinetes across the mosquito species tested (Figure 2a). For NF54, there was a tendency for fewer mature ookinetes in *An. gambiae* compared to *An. stephensi* and *An. coluzzii*, though this was not statistically significant (Figure 2a). A similar trend was observed with the ART-R ARN1G parasite line. For the Ugandan ART-R isolate PAT-023, significantly fewer mature ookinetes were observed in *An. stephensi* (Figure 2a) compared to *An. coluzzii* (p = 0.019). However, this significance level should be interpreted cautiously due to the number of comparisons made. The ART-R isolate 3815 consistently produced low numbers of mature ookinetes in all *Anopheles* species tested (Figure 2a).

**Figure 2.**
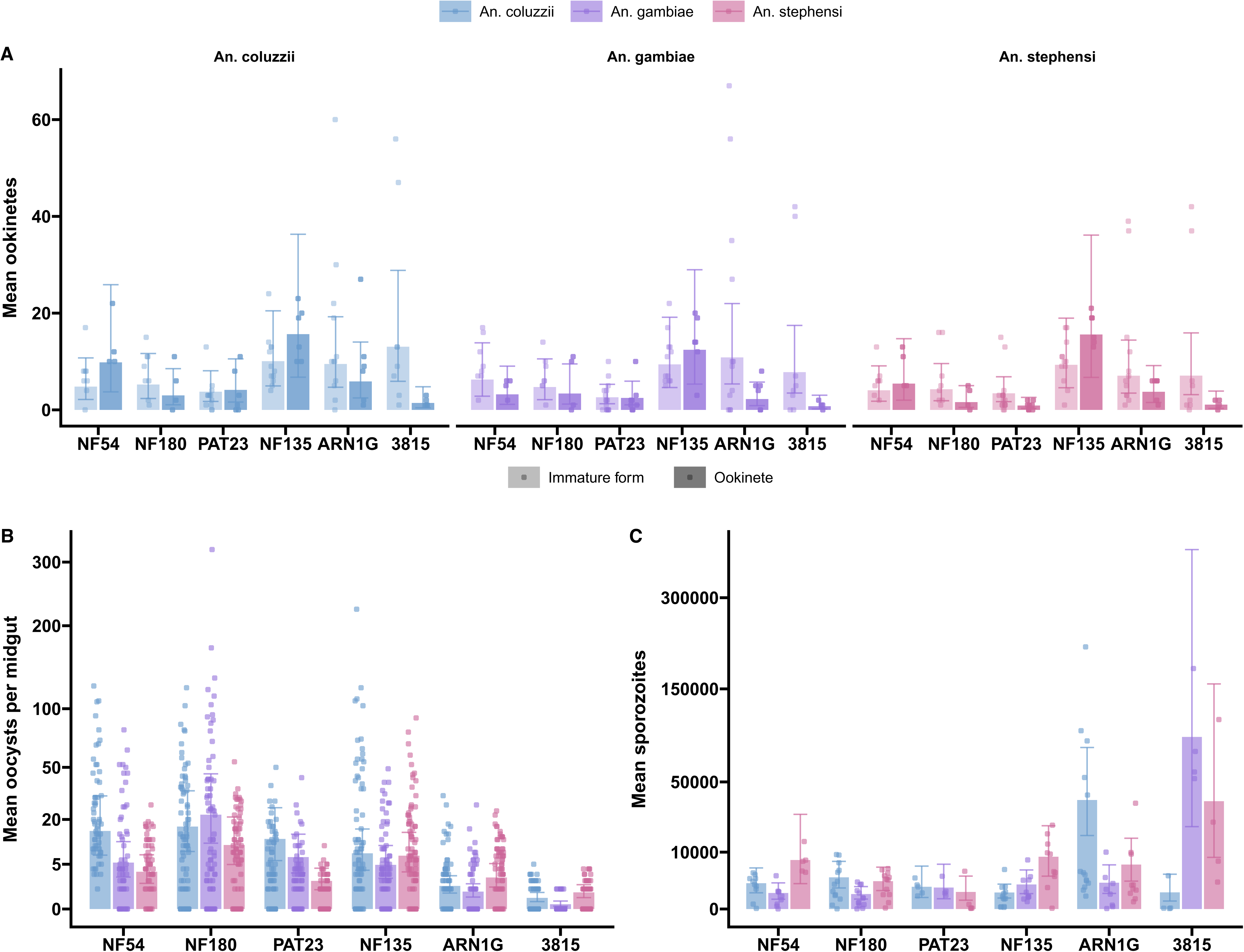
Infection of three mosquito species with the six different parasite lines. **A)** Ookinete counts 20 hours after blood meal. The ookinetes were stained with anti-Pfs25 488 conjugate and counted on a haemocytometer under a fluorescent microscope. Immature forms included rounded zygotes (or unfertilised females) and ookinetes with incomplete maturation (lighter shading bar), while mature forms were only the completely mature ookinetes (dark shaded bar). Dots represent a single count from an independent experiment and the error bars represent 95% confidence intervals. **B)** Average oocysts per midgut were dissected on day 7 post-bloodmeal. A total of 20 mosquitoes per group were dissected and oocysts were counted by mercurochrome staining. Dots represent the counted oocysts from a single midgut; error bars represent 95% confidence intervals. **C)** Salivary glands from individual mosquitoes were dissected and sporozoites quantified by qPCR. The dots represent the sporozoites per salivary gland from a single mosquito and the error bars represent the confidence intervals.

To confirm that mature ookinetes can establish mosquito infections, we quantified oocyst prevalence (Sup Figure 1) and density (Figure 2b) on day 7 post-bloodmeal. African parasite lines (NF54, NF180, and PAT-023) exhibited higher oocyst densities in traditional African mosquitoes (*An. gambiae s.s., An. coluzzii*) compared to *An. stephensi,* particularly for PAT-023 (p <0.0001; Sup Table 3). In contrast, NF135 showed consistent infection intensities across all tested mosquito species. For the Asian K13 mutant lines, oocysts densities in *An. stephensi* were slightly higher compared to *An. coluzzii* and *An. gambiae*. Specifically, ARN1G had higher densities in *An. stephensi* relative to *An. coluzzii* (p = 0.009) and *An. gambiae* (p < 0.0001), while 3815 showed similar trends (p = 0.007 compared to *An. coluzzii* and p<0.0001 compared to *An. gambiae*; Sup Table 3). The 3815 isolate exhibited low and highly variable infection levels, consistent with its reduced mature ookinete counts (Figures 2a and 2b). Interestingly, a trend towards higher sporozoite production was observed in the two K13 mutant lines (ARN1G and 3815), which persisted even after normalising for oocyst density within the same batch of mosquitoes. However, transmission results for the 3815 isolate remained highly variable (Sup Figure 2).

### Transmission-blocking effect of DHA

With all parasite lines being able to infect mosquitoes in the absence of drug pressure, we determined the transmission-reducing effect of DHA on mosquito infection intensity and prevalence. Mature gametocytes were exposed to both the physiologically relevant concentration of 700nM and a tenfold higher concentration of 7000nM for 48 hours before mosquito feeding with the drug not removed before mosquito feeding [32, 45]. Given the labour-intensive nature of these assays and the known permissiveness of *An. coluzzii* for the parasites tested herein, experiments were performed exclusively with this mosquito species. All isolates showed a consistent decrease in mosquito infection intensity and prevalence as DHA concentrations increased, independent of the K13 genotype and the parasite background (Figure 3, Sup Figure 3). At 7000nM DHA, sporadic infections were observed for the African isolates NF54, NF180, and PAT-023, but none of the Asian isolates infected mosquitoes at this concentration. When relative reductions in oocyst intensity were calculated, DHA exposure at 7000nM reduced oocyst intensities by >93% for all parasite isolates. There were no indications of reduced transmission-blocking efficacy of DHA for ART-R isolates.

**Figure 3.**
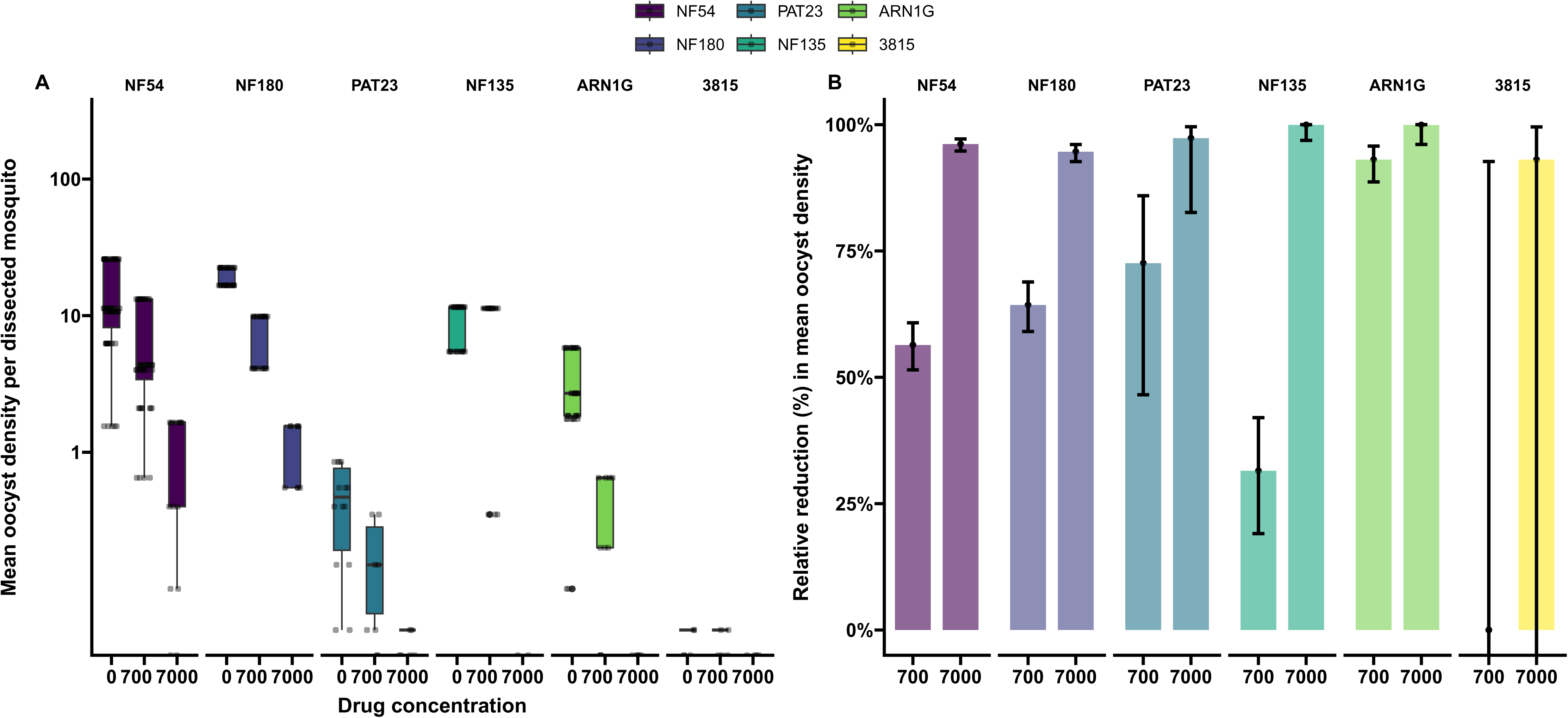
Transmission reduction in the presence of DHA. **A)** Oocyst densities observed upon exposing mature gametocytes to 700nM and 7000nM DHA prior to being fed to mosquitoes in a blood meal. The dots represent the average oocysts per infected mosquito from a single cage. **B)** Relative reductions in oocysts density compared the no drug control. All error bars represent the confidence intervals.

### Correlating resistance and transmission stages

Although the number of parasite isolates examined was modest, our data on asexual parasite survival under DHA pressure, combined with findings on sexual conversion and mosquito transmission, provide an opportunity to investigate whether ART-R confers a transmission advantage. We initially hypothesised that parasites with high levels of *in vitro* resistance to ART at the ring stage would show enhanced transmission to mosquitoes under drug pressure. However, no correlation was observed between parasite survival in the RSA and the reduction in oocyst density under DHA pressure (Figure 4a). In contrast with our initial hypothesis, we observed a weak negative correlation between parasite survival rates in the RSA and gametocyte conversion rates upon induction (Figure 4b).

**Figure 4.**
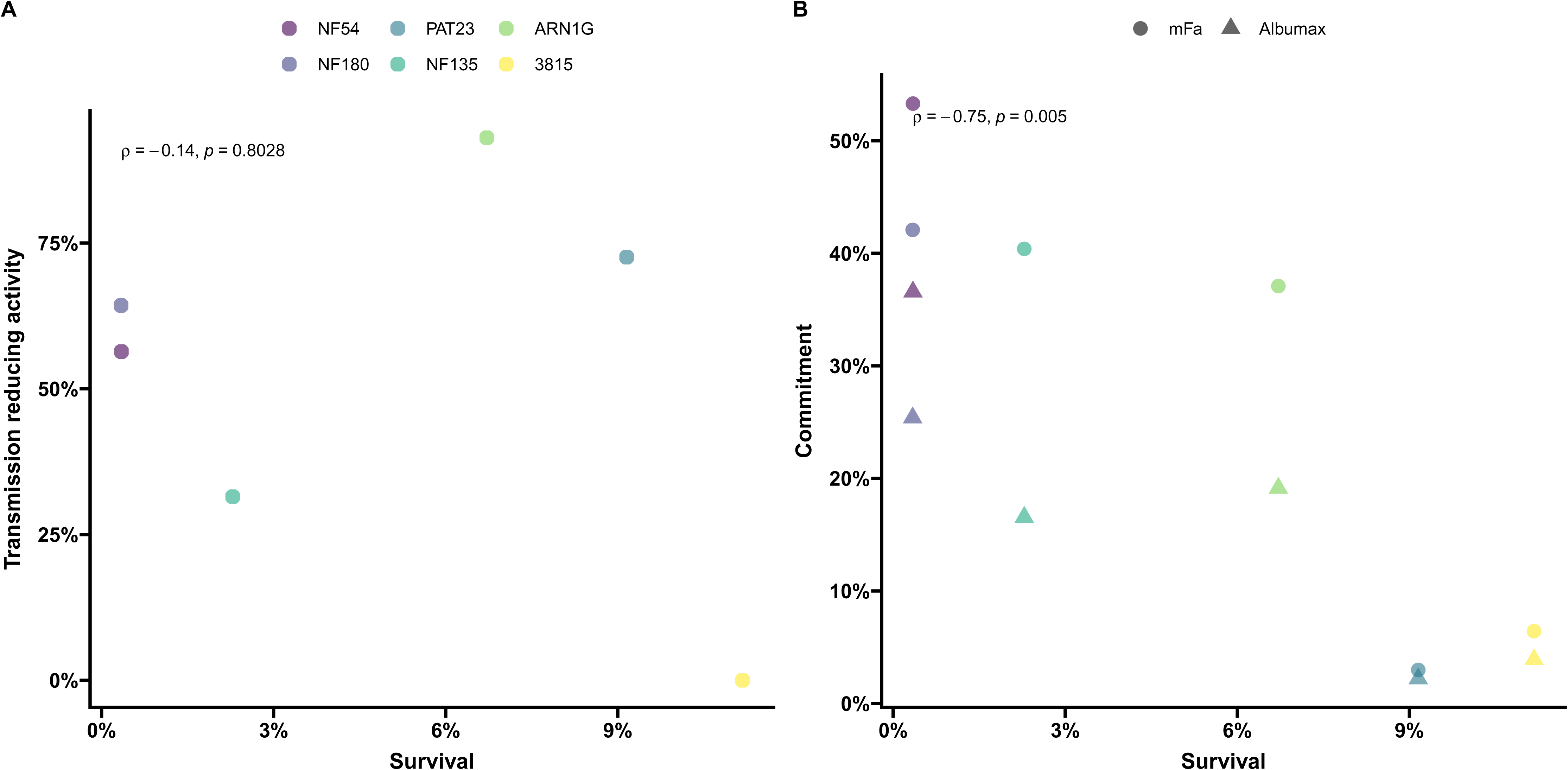
Impact of resistance on transmission and commitment. Transmission **A)** and conversion **B)** plotted against survival rates from the RSA for each parasite line. The underlying detailed data on *in vitro* parasite resistance in the RSA and gametocyte commitment are presented in Figure 1; data on transmission under DHA exposure are presented in Figure 3.

## Discussion

Understanding the intricacies of transmission fitness of parasites that survive ART-based treatment is important for developing strategies that aim to prevent or slow down the spread of resistance. Some reports have suggested that K13 mutant parasites may have higher intrinsic gametocyte production [4, 46], yet some *in vivo* evidence suggests following ART treatment in a Vietnam cohort, the conversion marker *ap2g* decreases – suggesting a decrease in gametocyte production linked to K13 mutations [47]. In contrast, an *in vitro* study observed increased survival of specifically male gametocytes of K13 mutants under DHA exposure [32]. We selected three parasite isolates from Southeast Asia and three from Sub-Saharan Africa, each with distinct K13 genotypes and genetic backgrounds. The Asian ART-R lines ARN1G and 3815 exhibited higher parasite survival in the RSA, consistent with previous studies [16, 32, 35]. PAT-023, a newly characterised Ugandan ART-R line that is wild-type for K13, demonstrated a high level of survival in the *in vitro* RSA. Interestingly, we also observed that NF135, a line previously associated with treatment failure following artemether-lumefantrine therapy *in vivo* [48], showed erratic and occasionally increased DHA survival rates.

This study focuses on the transmission potential of *P. falciparum* isolates, beginning with their commitment to sexual stages. Fatty acids play a key role in signalling the parasite to commit to sexual conversion [36, 37]; we used two methods targeting this signalling pathway to robustly compare sexual conversion rates. From all media conditions tested, minimal fatty acid media consistently produced the highest sexual conversion rates across all lines. Among the two ART-S parasite isolates, NF54 and NF135 consistently exhibited high conversion levels. The newly introduced NF180 line, which has not been characterised in detail previously, demonstrated high conversion rates, comparable to NF54 and higher than NF135. Among the ART-R parasite lines, PAT-023 displayed the lowest commitment to gametocyte production, while ARN1G showed commitment levels similar to NF135. The Cambodian 3815 line exhibited an inconsistent commitment to gametocyte production, with occasional high levels that decreased with prolonged culture, potentially reflecting epigenetic silencing during *in vitro* cultivation.

We observed no evidence of increased sexual conversion in the ART-R isolates in the absence of treatment. Of note, we did not explore the possible impact of DHA on conversion rates in these ART-R lines. An earlier study reported that the ART-S NF54 line was associated with higher sexual conversion when trophozoites were treated with DHA [49]. Future studies that investigate whether DHA-treated ART-R lines may give rise to an elevated conversion rate can be of value. Such studies would allow us to disentangle the contributions to the overall transmission competence; if this is linked to the increased conversion to or survival of the sexual stages after ART treatment or an inherent increased conversion rate in the K13 mutant lines.

While sexual conversion is a critical step for transmission, high commitment rates do not always correlate with high mosquito infectivity [25, 29]. The presence of mature ookinetes is a direct indicator of fertilisation in the mosquito environment. In our study, we fed the same gametocyte material to the three mosquito species. We did not normalize for gametocyte density prior to feeding; instead we had a uniform starting parasite concentration prior to gametocyte induction. We are thus able to compare the composite product of gametocyte production and infectivity between lines. When examining the impact of individual mutations on transmission competence in matched genetic backgrounds, such normalization at different points in the transmission cycle may be beneficial. We reproducibly observed low transmission potential of the 3815 line, which consistently formed very few ookinetes in all species and exhibited low and sporadic oocyst infections. We also observed a tendency for African parasites to fare better in the African mosquitoes, illustrated by both the numbers of mature ookinetes and oocysts densities. Although our sample size is too small for a comprehensive analysis of the underlying biological mechanisms, one could speculate that genes influencing the parasite’s ability to evade the mosquito immune system may play a role. Pfs47 is essential for *P. falciparum* infection in *An. gambiae* but not for *An. stephensi* [50, 51]. Contrary to this observation for African parasite lines, all three Southeast Asian parasite lines were able to infect Asian and African mosquito vectors. Interestingly ARN1G achieved slightly higher ookinete numbers in *An. coluzzii* but this did not translate into higher oocyst densities. Similar to our observations on sexual conversion rates, we found no significant transmission advantage for ART-R parasite lines with reduced ART susceptibility in the RSA.

To confirm the ability of the parasites to complete sporogony, we quantified sporozoite numbers in mosquito salivary glands. Previous studies have suggested that K13 mutant lines might produce larger oocysts [32], potentially indicating increased sporozoite production. While we did not observe a clear increase in oocyst size (data not shown) whilst sporozoite production varied between parasite lines. The two Asian ART-R lines, each with a unique K13 mutation, exhibited significantly higher sporozoite loads in at least one African mosquito species. This effect was most pronounced for ARN1G that showed significantly higher sporozoite numbers in *An. coluzzii*, even after adjusting for oocyst density within the same mosquito batch. Further studies are needed to confirm these findings and to explore the underlying biological mechanisms. However, our results are consistent with the suggestion that the K13 mutation may lead to larger oocysts [32], which could lead to increased sporozoite production.

Lastly, we examined the impact of DHA exposure on transmission efficiency, hypothesising that parasite lines with increased survival rate of asexual parasites under DHA exposure may show similar survival advantages when gametocytes are exposed. Previous work has suggested that male gametocytes with K13 mutations may have partial protection against DHA [32, 52]. Using *An. coluzzii,* we tested two DHA concentrations and found no evidence that DHA was less effective against mature gametocytes and subsequently reducing transmission in ART-R parasites. Immature gametocytes are known to be more susceptible to artemisinins than mature gametocytes [32, 53]. Since we did not examine the impact of DHA on immature gametocytes, we cannot rule out preferential survival of immature ART-R gametocytes. Even at the highest DHA concentrations, both ART-S and ART-R African parasite lines occasionally infected mosquitoes. This observation is consistent with previous *ex vivo* experiments [45] and aligns with clinical studies showing that treatment with DHA-piperaquine fails to fully prevent transmission in the first weeks after treatment [54].

In this study, we examined a limited number of ART-S and ART-R parasite lines across three mosquito species. We observed no consistent evidence of increased gametocyte production or increased mosquito infectivity. While the transmission-reducing effect of DHA was imperfect, its efficacy was not reduced in gametocyte-producing lines with partial resistance to ARTs. To fully uncover the discrete impacts of individual K13 mutations on gametocyte production, infectivity and gametocyte resistance to ARTs, future studies using genetically engineered lines in controlled isogenic backgrounds are required.

## Ethics declarations

Experiments with *in vitro* cultured parasites and *Anopheles* mosquitoes at Radboud University Medical Center were conducted following approval from the Radboud University Experimental Animal Ethical Committee (RUDEC 2009-019, RUDEC 2009-225). PAT-023 was obtained with patient informed consent for sample collection and for its future use. The study protocol that this collection was part of was approved by the Makerere University School of Biomedical Sciences Research and Ethics Committee (SBS-363), the Uganda National Council for Science and Technology, and the University of California, San Francisco Committee on Human Research [55].

## Acknowledgements and financial support

We would like to thank Jolanda Klaassen, Laura Pelser-Posthumus, Astrid Pouwelsen and Jacqueline Kuhnen for breeding of mosquitoes and handling of the infected mosquitoes. We are grateful for the support of Marga de Vegte-Bolmer in starting parasite culture with the 3815 line. DAF gratefully acknowledges funding support from the NIH (R01 AI109023). SM acknowledges the support from the Human Frontier Science Program Long-term Fellowship (LT000976/2016-L) and funding from United States National Institutes of Health NIAID grant R01 AI182318 (PI: SM). This work was supported by a fellowship from the Netherlands Organization for Scientific Research (Vici fellowship NWO 09150182210039).

## Supplemental Tables

**Sup Table 1.** RSA comparisons.

| Reference parasite line | Parasite line | Ratio (CI) |
| --- | --- | --- |
| NF54 | NF180 | 0.99 (0.25, 3.85); p=0.9742 |
| NF54 | PAT-023 | 33.34 (10, 100); p<0.0001 |
| NF54 | NF135 | 6.67 (1.49, 33.34); p=0.0023 |
| NF54 | ARN1G | 20.00 (6.25, 100); p<0.0001 |
| NF54 | 3815 | 33.34 (12.5, 100); p<0.0001 |
| NF135 | PAT-023 | 4.31 (1.20, 15.49); p=0.0057 |
| NF135 | ARN1G | 3.13 (0.81, 12.5); p=0.0636 |
| NF135 | 3815 | 5.26 (1.51, 20); p=0.0014 |

**Sup Table 2.** Sexual conversion comparisons.

| Parasite line | Reference media | Media | Ratio (CI) |
| --- | --- | --- | --- |
| NF54 | serum | mFa | 114.14 (31.57, 412.62); p<0.0001 |
| NF54 | serum | Albumax | 57.67 (15.77, 210.87); p<0.0001 |
| NF54 | mFa | Albumax | 0.51 ( 0.16, 1.60); p=0.1516 |
| NF180 | serum | mFa | 46.96 (14.23, 154.93); p<0.0001 |
| NF180 | serum | Albumax | 21.99 ( 6.47, 74.73); p<0.0001 |
| NF180 | mFa | Albumax | 0.47 ( 0.16, 1.39); p=0.0934 |
| PAT-023 | serum | mFa | 3.95 ( 1.06, 14.70); p=0.0374 |
| PAT-023 | serum | Albumax | 2.90 ( 0.78, 10.79); p=0.1027 |
| PAT-023 | mFa | Albumax | 0.73 ( 0.20, 2.74); p=0.5691 |
| NF135 | serum | mFa | 92.75 (25.51, 337.27); p<0.0001 |
| NF135 | serum | Albumax | 27.16 ( 6.95, 106.13); p<0.0001 |
| NF135 | mFa | Albumax | 0.29 ( 0.09, 1.01); p=0.0172 |
| ARN1G | serum | mFa | 75.74 (22.86, 250.88); p<0.0001 |
| ARN1G | serum | Albumax | 30.40 ( 8.75, 105.61); p<0.0001 |
| ARN1G | mFa | Albumax | 0.40 ( 0.13, 1.24); p=0.0511 |
| 3815 | serum | mFa | 7.27 ( 1.97, 26.84); p=0.0011 |
| 3815 | serum | Albumax | 4.30 ( 1.16, 16.00); p=0.0162 |
| 3815 | mFa | Albumax | 0.59 ( 0.16, 2.19); p=0.3311 |

552 mFa, minimal fatty acid.

**Sup Table 3.**
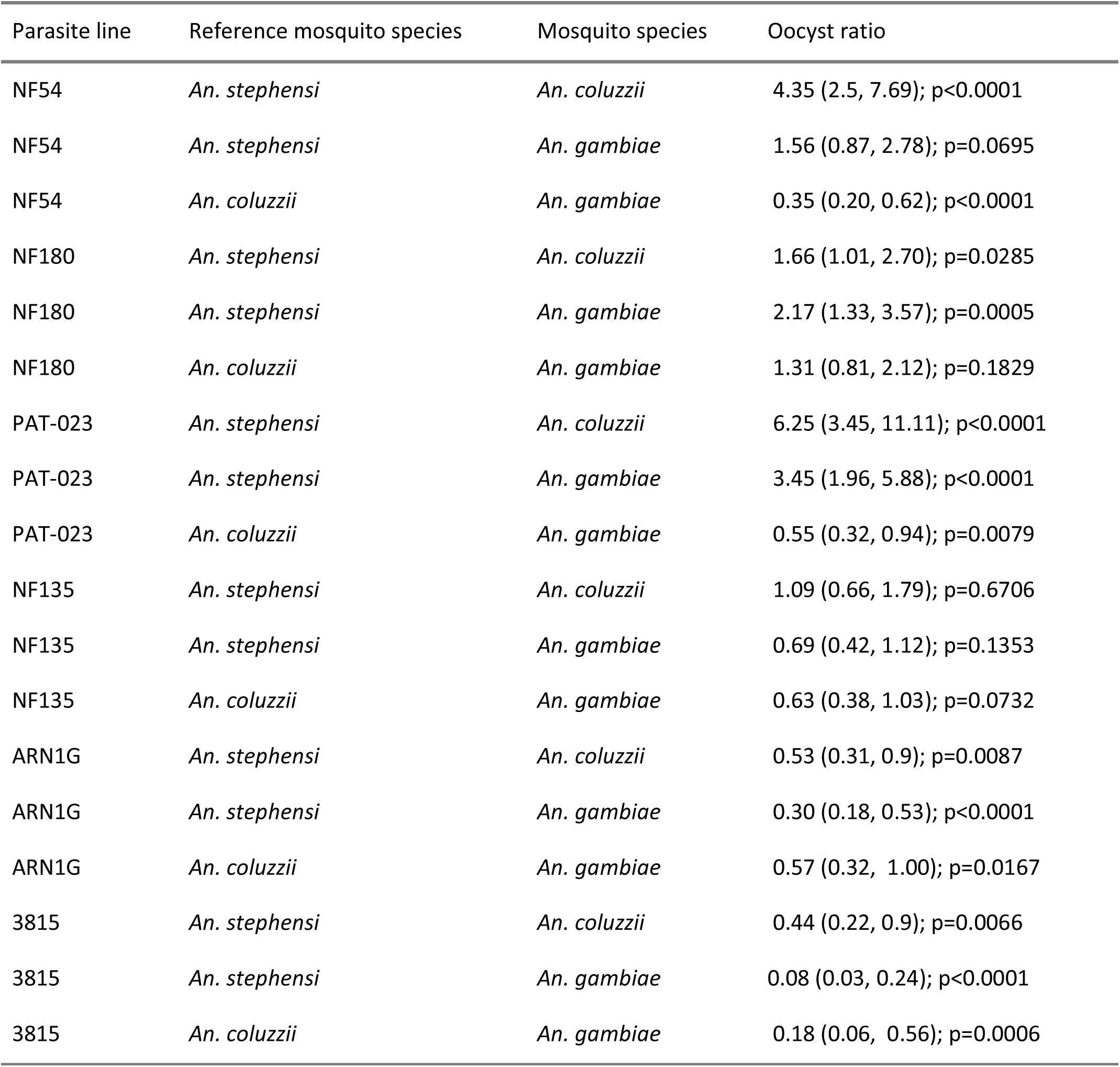
Oocyst comparisons.

**Sup Table 4.**
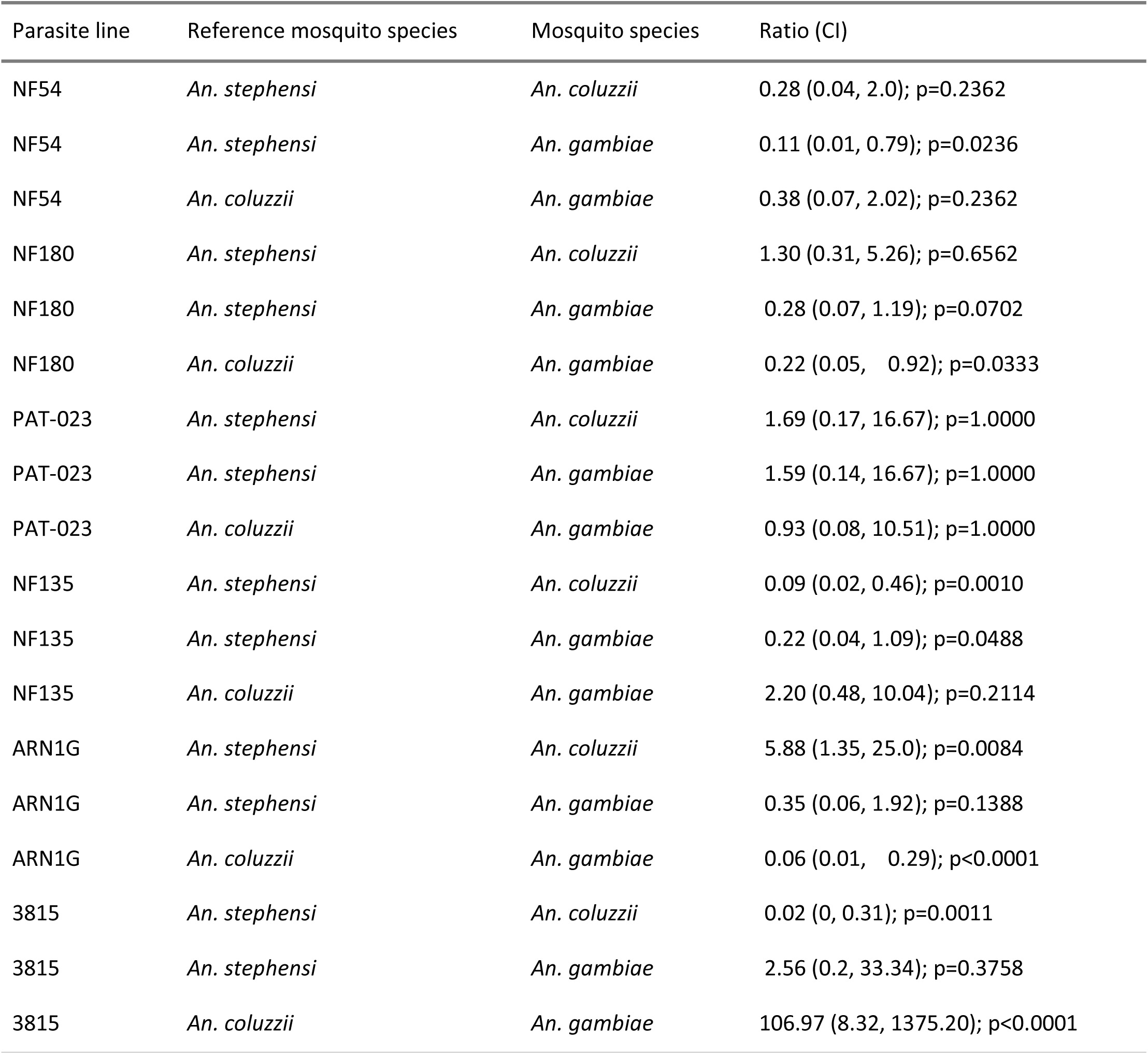
Sporozoite comparisons.

| Parasite line | Reference mosquito species | Mosquito species | Ratio (CI) |
| --- | --- | --- | --- |
| NF54 | *An. stephensi* | *An. coluzzii* | 0.28 (0.04, 2.0); p=0.2362 |
| NF54 | *An. stephensi* | *An. gambiae* | 0.11 (0.01, 0.79); p=0.0236 |
| NF54 | *An. coluzzii* | *An. gambiae* | 0.38 (0.07, 2.02); p=0.2362 |
| NF180 | *An. stephensi* | *An. coluzzii* | 1.30 (0.31, 5.26); p=0.6562 |
| NF180 | *An. stephensi* | *An. gambiae* | 0.28 (0.07, 1.19); p=0.0702 |
| NF180 | *An. coluzzii* | *An. gambiae* | 0.22 (0.05, 0.92); p=0.0333 |
| PAT-023 | *An. stephensi* | *An. coluzzii* | 1.69 (0.17, 16.67); p=1.0000 |
| PAT-023 | *An. stephensi* | *An. gambiae* | 1.59 (0.14, 16.67); p=1.0000 |
| PAT-023 | *An. coluzzii* | *An. gambiae* | 0.93 (0.08, 10.51); p=1.0000 |
| NF135 | *An. stephensi* | *An. coluzzii* | 0.09 (0.02, 0.46); p=0.0010 |
| NF135 | *An. stephensi* | *An. gambiae* | 0.22 (0.04, 1.09); p=0.0488 |
| NF135 | *An. coluzzii* | *An. gambiae* | 2.20 (0.48, 10.04); p=0.2114 |
| ARN1G | *An. stephensi* | *An. coluzzii* | 5.88 (1.35, 25.0); p=0.0084 |
| ARN1G | *An. stephensi* | *An. gambiae* | 0.35 (0.06, 1.92); p=0.1388 |
| ARN1G | *An. coluzzii* | *An. gambiae* | 0.06 (0.01, 0.29); p<0.0001 |
| 3815 | *An. stephensi* | *An. coluzzii* | 0.02 (0, 0.31); p=0.0011 |
| 3815 | *An. stephensi* | *An. gambiae* | 2.56 (0.2, 33.34); p=0.3758 |
| 3815 | *An. coluzzii* | *An. gambiae* | 106.97 (8.32, 1375.20); p<0.0001 |

**Sup Table 5.** Sporozoite per oocyst comparisons.

| Parasite line | Reference mosquito species | Mosquito species | Sporozoite per oocyst ratio |
| --- | --- | --- | --- |
| NF54 | *An. stephensi* | *An. coluzzii* | 0.06 (0.01, 0.50); p=0.0044 |
| NF54 | *An. stephensi* | *An. gambiae* | 0.07 (0.01, 0.56); p=0.0047 |
| NF54 | *An. gambiae* | *An. coluzzii* | 0.92 (0.16, 5.33); p=0.9092 |
| NF180 | *An. stephensi* | *An. coluzzii* | 0.78 (0.17, 3.51); p=0.6919 |
| NF180 | *An. stephensi* | *An. gambiae* | 0.13 (0.03, 0.60); p=0.0042 |
| NF180 | *An. gambiae* | *An. coluzzii* | 5.89 (1.32, 26.36); p=0.0096 |
| PAT-023 | *An. stephensi* | *An. coluzzii* | 0.27 (0.03, 2.88); p=0.5525 |
| PAT-023 | *An. stephensi* | *An. gambiae* | 0.46 (0.04, 5.59); p=0.9104 |
| PAT-023 | *An. gambiae* | *An. coluzzii* | 0.59 (0.05, 7.06); p=0.9104 |
| NF135 | *An. stephensi* | *An. coluzzii* | 0.09 (0.02, 0.45); p=0.0012 |
| NF135 | *An. stephensi* | *An. gambiae* | 0.32 (0.06, 1.74); p=0.1198 |
| NF135 | *An. gambiae* | *An. coluzzii* | 0.29 (0.06, 1.42); p=0.1198 |
| ARN1G | *An. stephensi* | *An. coluzzii* | 11.30 (2.31, 55.28); p=0.0009 |
| ARN1G | *An. stephensi* | *An. gambiae* | 1.15 (0.19, 6.97); p=0.8499 |
| ARN1G | *An. gambiae* | *An. coluzzii* | 9.81 (1.79, 53.69); p=0.0028 |
| 3815 | *An. stephensi* | *An. coluzzii* | 0.05 (0.00, 0.76); p=0.0084 |
| 3815 | *An. stephensi* | *An. gambiae* | 31.40 (1.95, 504.25); p=0.0062 |
| 3815 | *An. gambiae* | *An. coluzzii* | 0.00 (0.00, 0.03); p<0.0001 |

