## Supplementary figures and images for "Gametocyte production and transmission fitness of African and Asian *Plasmodium falciparum* isolates with differential susceptibility to artemisinins"

### Supplemental Figure 1

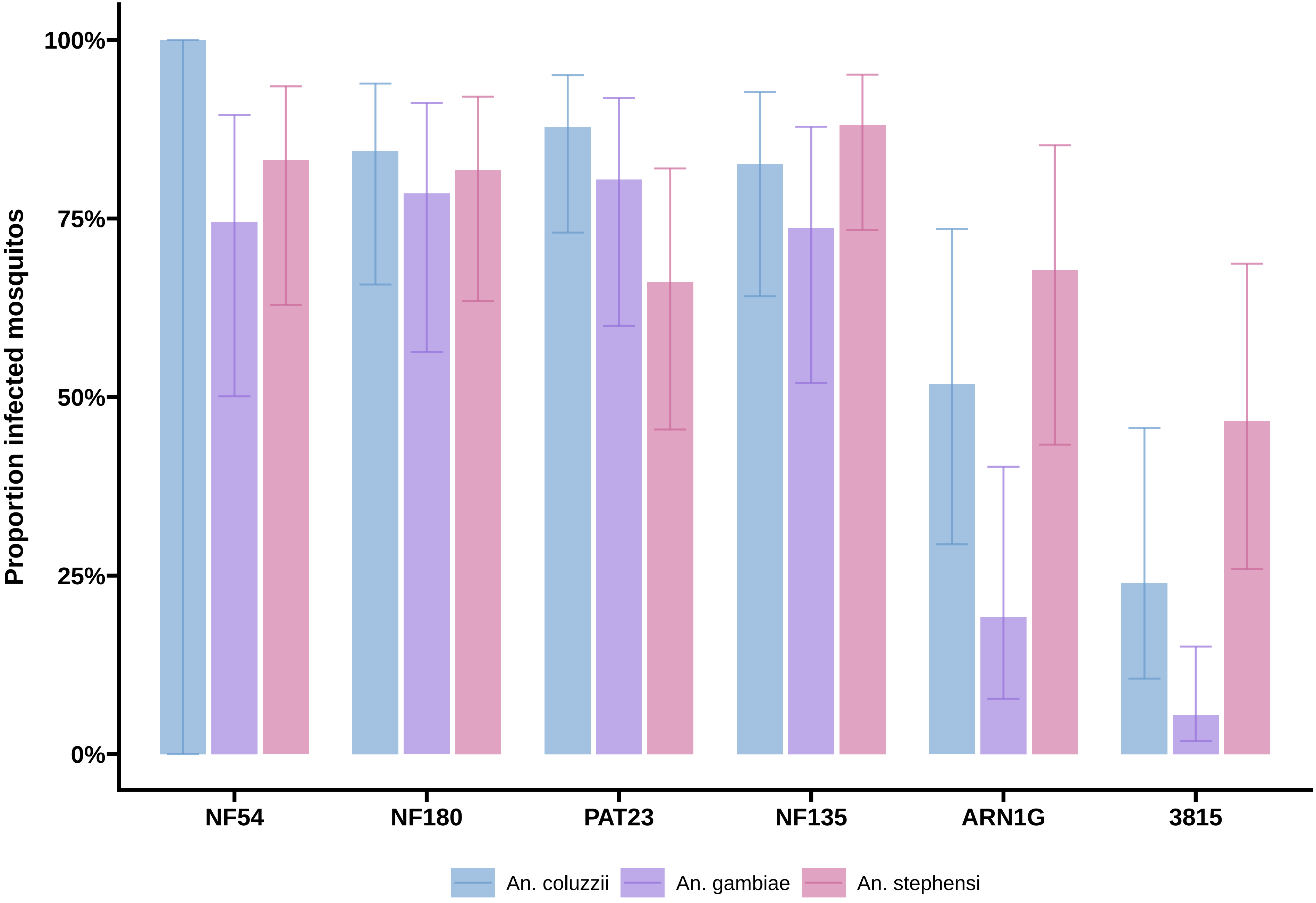

### Supplemental Figure 2

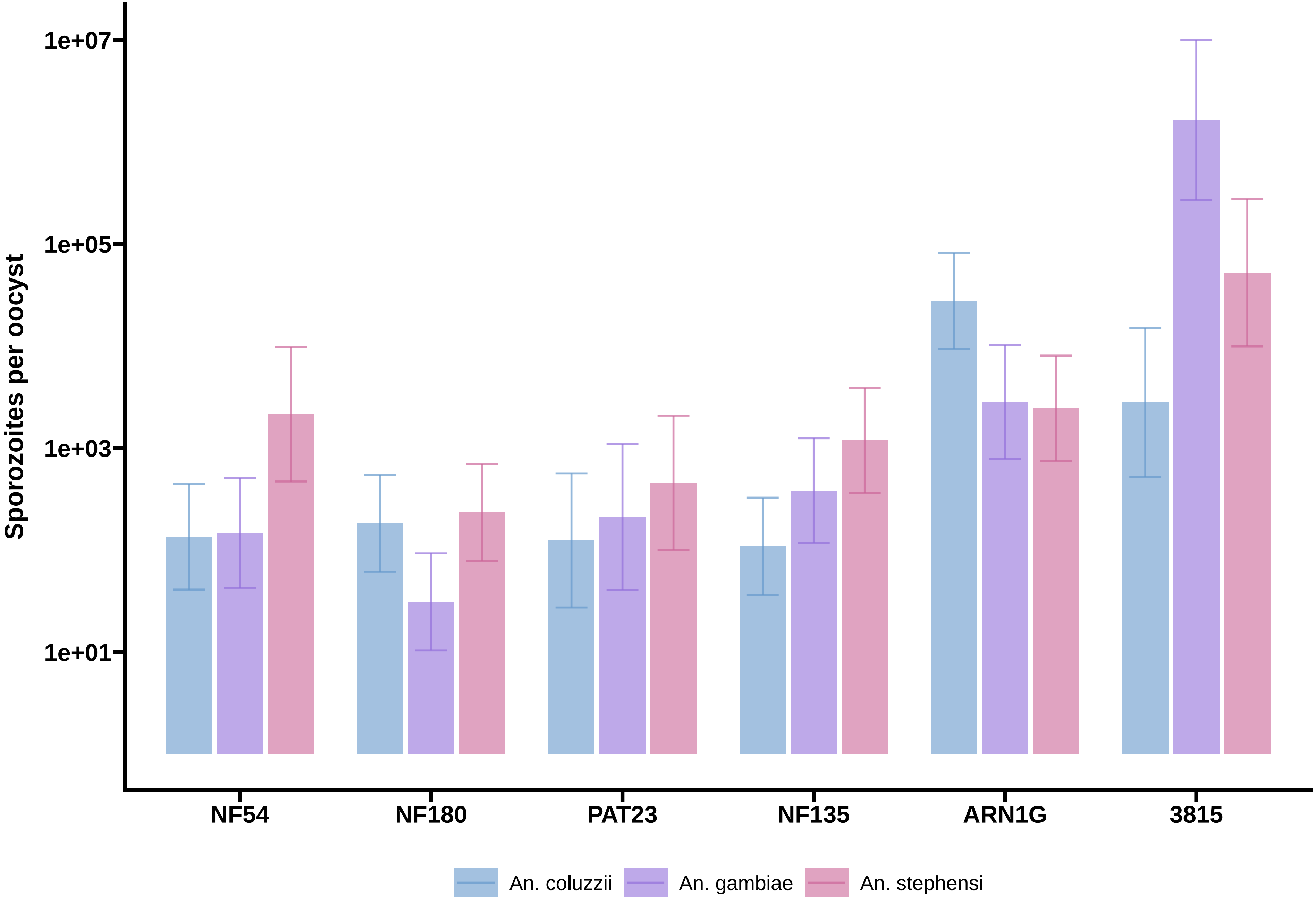

### Supplemental Figure 3

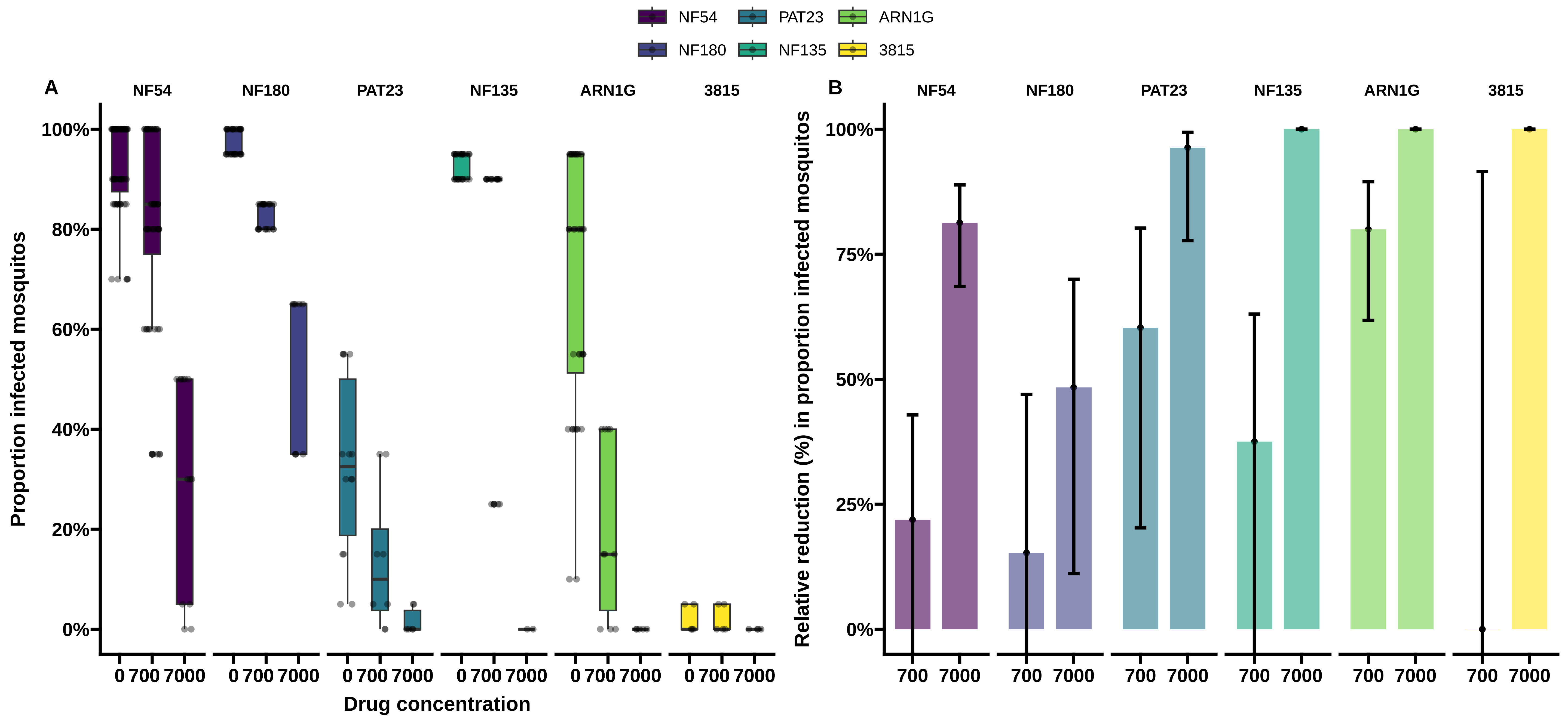
